# Subpopulation Codes Permit Information Modulation Across Cortical States

**DOI:** 10.1101/2022.09.28.509815

**Authors:** Matthew P. Getz, Chengcheng Huang, Brent Doiron

## Abstract

Cortical state is modulated by myriad cognitive and physiological mechanisms. Yet it is still unclear how changes in cortical state relate to changes in neuronal processing. Previous studies have reported state dependent changes in response gain or population-wide shared variability, motivated by the fact that both are important determinants of the performance of any population code. However, if the state-conditioned cortical regime is well-captured by a linear input-output response (as is often the case), then the linear Fisher information (FI) about a stimulus available to a decoder is invariant to state changes. In this study we show that by contrast, when one restricts a decoder to a subset of a cortical population, information within the subpopulation can increase through a modulation of cortical state. A clear example of such a subpopulation code is one in which decoders only receive projections from excitatory cells in a recurrent excitatory/inhibitory (*E/I*) network. We demonstrate the counterintuitive fact that when decoding only from *E* cells, it is exclusively the *I* cell response gain and connectivity which govern how information changes. Additionally, we propose a parametrically simplified approach to studying the effect of state change on subpopulation codes. Our results reveal the importance of inhibitory circuitry in modulating information flow in recurrent cortical networks, and establish a framework in which to develop deeper mechanistic insight into the impact of cortical state changes on information processing in these circuits.

## Introduction

Cortical circuits encode information about stimulus or action variables in their population activity [47]. These circuits then act on other cortical, musculoskeletal or endocrine systems to drive behavior. Studying cortical processing as an information transmission problem is a useful step toward relating neuronal activity and behavioral responses [39, 42, 27]. Theories of population coding require an understanding of both the sensitivity of trial-averaged stimulus tuning [49] and the structure of population-wide trial-to-trial variability [1, 3, 26]. Any changes in these two measures of neuronal response must be considered in order to evaluate whether the stimulus information available to a decoder of the population has increased or decreased. Such analysis has helped interpret experimental observations; for instance attention, well known to improve behavior in complex visual tasks [40], has been found to increase firing rates and neural selectivity while decreasing pairwise noise correlations along the visual pathway [34, 33, 5, 45, 41]. These effects are therefore taken to coincide with an attention-mediated improvement in information processing [54].

Biologically-motivated models of neuronal circuits define a network’s state as the dynamical regime in which it is operating [19]. Top-down modulatory processes engaged during shifts in attention or arousal induce changes in the state of a cortical network through processes such as neuromodulation, feedback projections, and synaptic rearrangement [9, 35, 17, 31]. Any shift in network state will affect how a network responds to stimuli, observed as a shift in trial averaged tuning as well as response variability. Uncovering network mechanisms which enable cognitive processes like attention to improve behavioral performance must therefore take network state into account. Yet prior studies of information processing have focused on parametric models that are agnostic to underlying circuit mechanisms, so that any shift in tuning is made without considering a concomitant shift in variability (or vice versa) [49, 2, 3, 24, 1, 21]. There thus remains a gap in our mechanistic understanding of the way in which information changes as a function of cortical state.

In this study we explore how changes in cortical state affect information processing in neural circuits. We find that in any circuit, the information available to a linear decoder is invariant to network state when the decoder reads out from the entire population [23]. However, in many cases only a subset of the neurons project to a given decoder: inhibitory neurons have predominantly local projections [55] while excitatory neurons are often subdivided based on their outward projections [50, 57]. From the vantage of information processing we label this a *subpopulation code*. We show that when a network’s state is modulated then the information available to a linear decoder of a subpopulation code is malleable. Intriguingly, we also find that the information encoded in a subpopulation does not explicitly depend on the activity and connectivity within the subpopulation. Rather, it is only those neurons within the circuit which connect *to* the projection cells that directly shape information flow. We thereby demonstrate that it may not be possible to draw significant conclusions on state dependent cortical processing without a more complete view of the network in question. Towards this end, we provide a framework for a circuit dissection that exposes how modulations of cortical state impact information processing in neural circuits.

## Results

There are a vast array of mechanisms through which the brain modulates cortical network state. Two of those commonly studied, neuromodulators and feedback projections, can both uniquely affect circuit dynamics, from cellular excitability to transient synaptic weight changes [31, 17]. The neuronal correlates of these state changes are shifts in the firing rate, neuronal sensitivity, and correlations of populations of neurons. In relating these changes in neural response statistics to behavior we are really asking how a modulus influences cortical processing. Fisher information (FI) provides a means of addressing this question as it measures a decoder’s ability to discriminate between two stimuli. It has been argued that in the regime in which sensory cortex lives, linear FI is equivalent to FI (and this likely extends to other areas of cortex as well, where linear decoders have best fit the relationship of neural activity to behavior) [43, 26]. We therefore focus on linear FI in this work, which is given by

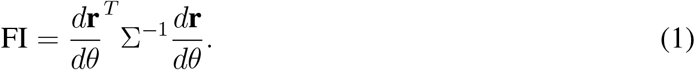

Here 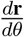 is the population response gain (change in rate for a change in a stimulus parameter *θ*) and Σ is the covariance matrix of the population response.

Now consider a recurrently coupled network of excitatory (*E*) and inhibitory (*I*) populations (Figure 1a) with dynamics described by

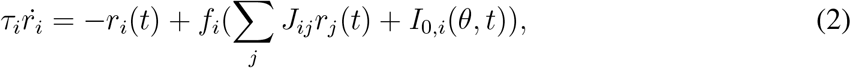

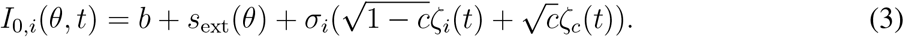

**Figure 1:**
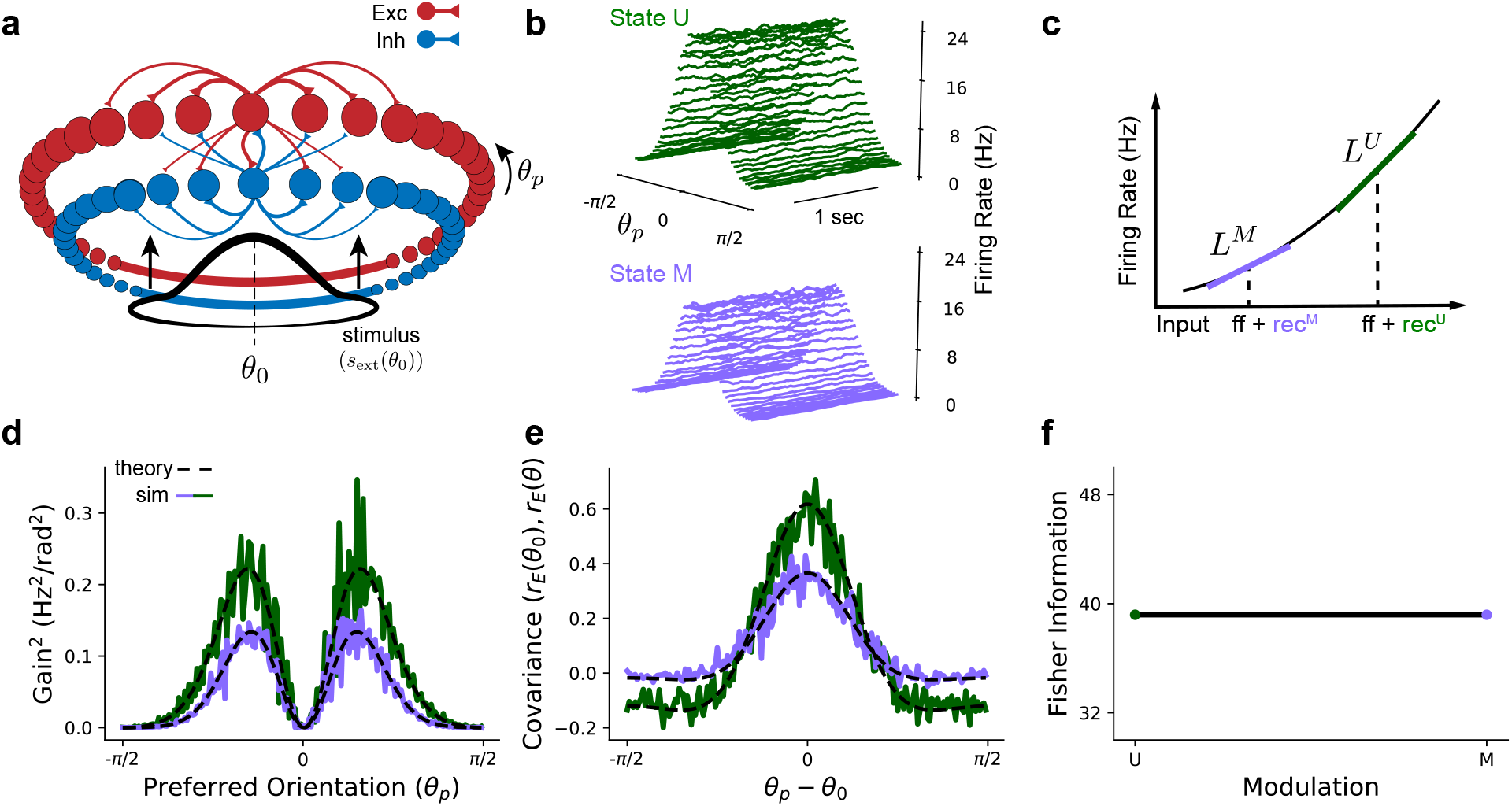
Changes in network activity do not imply changes in information. **a** Network schematic. Units are arranged on a ring indexed by *θ* where position on the ring corresponds to that unit’s preferred value of *θ*. Red units are excitatory and blue are inhibitory. Size of connection line indicates strength of connection. Not all connections are shown. **b**. Firing rates for the excitatory network units across time in the unmodulated (U) and modulated (M) states. **c**. Illustration of the effect of state changes on the input/output response for a single unit in the excitatory population before (green) and after (purple) modulation. **d**. Fit of the squared gain across the *E* population in a nonlinear model (solid lines) with a linear theory (black dashed lines). **e**. Fit of integrated cross-covariance relative to a single *E* unit (with preferred orientation *θ*_0_) with a linear theory. Colors as in (b). **f**. Plot of Fisher information as a function of modulation.

Here *i* is the index of a unit in the *E* or *I* population, *b* is baseline input for all cells (potentially heterogeneous but assumed constant for simplicity), *s*_ext_(*θ*) is an external drive that depends on the stimulus parameter *θ, J*_*ij*_ is the connection weight from unit *j* to unit *i, r*_*i*_ is the firing rate of unit *i* in the network, *f*_*i*_ is the transfer function of unit *i, ζ*_*i*_ is noise private to unit *i, ζ*_*c*_ is noise common to all units in the network and *c* scales the amount of shared variability (Eq. 14). We restrict our analysis to steady-state responses in which for sufficiently small input noise, the responses of the firing rate vector **r** can be linearized around the operating point (Figure 1c) [6]. Equation 2 then becomes:

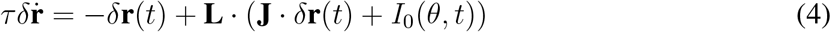

where *δ***r** are the dynamics of the response around the steady-state solution and **L** is the linearization matrix about the steady state rate (Eq. 13; [7]).

In the context of this system, a network modulation is a change in the steady-state rate without a change in stimulus input (*s*_ext_). We therefore define a modulation as a perturbation which changes the operating point of a network for a fixed stimulus (illustrated in Figure 1c). The space of perturbations is thus restricted to the space of network parameters (see also Figure 2a); some aspect of the network is changing such that we must relinearize about a new steady-state response. In this study we will consider moduli of two types: additive changes to the top-down input current (representative of cortical feedback [25]) or transient synaptic weight changes (e.g. neuromodulator-induced synaptic plasticity [31, 54]).

**Figure 2:**
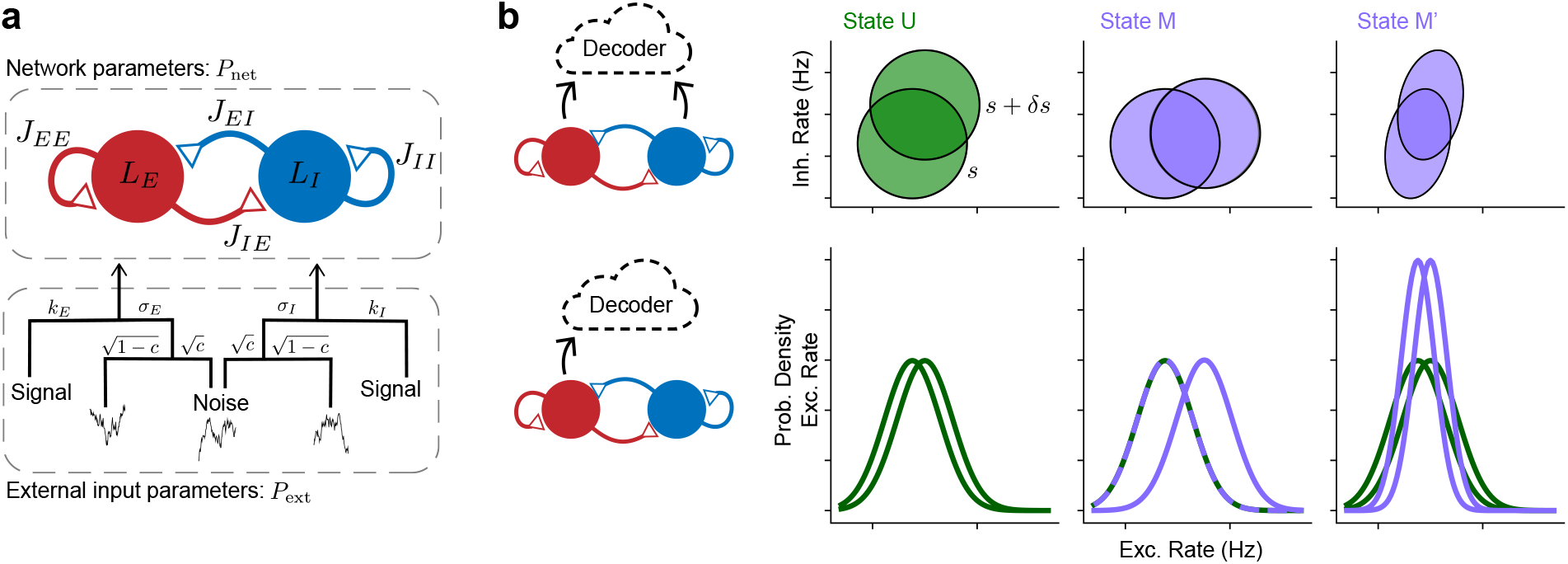
Projection to lower dimensions allows for improved discrimination with modulation. **a**. Illustration of *E/I* circuit parameters, partitioned into external input parameters (*P*_ext_; not affected by modulation) and network parameters (*P*_net_; subject to change through modulation). **b**. (Top row) 95th percentile distributions of idealized steady state rates at contrasts *s* and *s* + *δs* for unmodulated (green) and modulated (purple) networks. A decoder reading out from the full network (top left) has access to the joint *r*_*E*_*/r*_*I*_ distribution. Modulation of a linear model does not change discriminability in high dimensions because the overlap (error) between the joint *r*_*E*_*/r*_*I*_ distributions over *s* and *s* + *δs* does not change from U to M or M’. (Bottom row) Projection of the joint *r*_*E*_*/r*_*I*_ distribution onto the *r*_*E*_ axis is the same as a decoder restricted to observing only the *E* population (bottom left). The state changes in the *r*_*E*_*/r*_*I*_ space permit increased discriminability in the *E* population due to decreased overlap of the distributions.

Previous modeling studies have captured the effects of attention on the response statistics of cortical circuits with a change in the top-down input drive to the network [23, 13, 20]. We introduced a similar top-down drive to a model recurrent *E/I* network to induce a state change (see Methods: Ring network). This modulus resulted in significant changes to the network response to a fixed stimulus: by contrast to the unmodulated network (state U), in the modulated network (state M) firing rates decreased across the excitatory population (Figure 1b) resulting in a decrease in the squared gain of the population (Figure 1d) and decorrelation across *E* cells (Figure 1e). Given that these statistics define FI, we expect these dramatic changes to the network’s response statistics to affect information flow in this circuit. What we instead observed is that FI is invariant to this modulation (Figure 1f). We show in the next section (Eq. 6) that this is true for *all* possible moduli under our definition. This not only apparently contradicts the necessity that behavioral improvements would be reflected in enhanced information processing in cortex but also questions the effect of these neural response statistics on cortical processes more generally.

### Modulation can improve information flow in subpopulation codes

In order to develop intuition for the invariance of FI to modulation we turn to an *E/I* network with a single *E* and single *I* unit (Figure 2a). The responses *r*_*E*_ and *r*_*I*_ to an input stimulus *s* and a perturbation *s* + *δs* can be described as a joint probability distribution over the rates (the top row of Figure 2b). When a modulation is applied, the shape and center of the response distributions for *s* and *s* + *δs* both vary in *E*-*I* firing rate space (Figure 2b, purple ellipses). For instance, modulation could induce a rotation of the distributions in *r*_*E*_-*r*_*I*_ space (Figure 2b, state M), or change the correlations between *r*_*E*_ and *r*_*I*_ (Figure 2b, state M’). Overall discriminability between *s* and *s* + *δs* does not change in either of these scenarios, however, since the total overlap of the distributions does not change. Hence the decoding error is constant across network state which is consistent with the result of the full network model described above (Figure 1f).

This would seem to suggest that modulation of cortical circuits is functionally insignificant, and that optimization for sensory discrimination should act on other mechanisms, such as feedforward thalamic inputs. However until now, we have made a tacit assumption that all neurons encode the stimulus variable. Yet from the standpoint of a downstream cortical area only those neurons which project to it will matter in the readout. In particular, we know from anatomical studies that excitatory (pyramidal) neurons are the dominant projection cells in cortex. Hence information read out of a population by downstream areas must be predominantly conveyed through excitatory pathways [46]. Since our decoder is functionally equivalent to a downstream region decoding an upstream signal, we therefore reconsider the same problem by decoding from the excitatory population alone. This corresponds to a projection of the joint *r*_*E*_*/r*_*I*_ firing rate distribution onto the *r*_*E*_ axis (Figure 2b, bottom row). Similarly to the joint distributions, the distributions of excitatory firing rates can change significantly with state as well. Critically, we observe a decrease in the overlap of the *r*_*E*_ distributions with modulation (decrease in the error), indicating an increase in the discriminability in the *E* population following each modulation. Thus, while FI for the full network is invariant to state modulation, we see that FI restricted to the *E* population alone can change with state.

To formalize the arguments above in terms of FI, we define external stimuli for this *E/I* network as *s*_ext,*E*_ = *k*_*E*_*s, s*_ext,*I*_ = *k*_*I*_*s* in equation 3, where *k*_*α*_ is a sensitivity term which scales the size of a feedforward stimulus drive to population *α* ∈ {*E, I*}. Then the linear FI for the full *E/I* network is given by [23]:

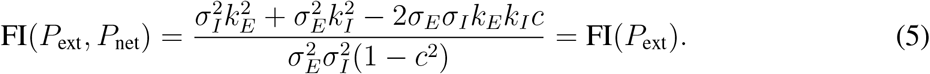

This equation is independent of all network parameters (*P*_net_), depending only on the external input parameters, *P*_ext_ (Figure 2a). Since we have defined a modulation to only affect the network state (and consequently only impact network parameters *P*_net_), FI for the full population must be invariant to modulation.

This result is true in general for any network size; linear FI reduces to a simple form which depends only on the input gain and input covariance since the output gain and output covariance depend on the network linearization in the same way which subsequently cancels (see Methods). Thus, in a linearized system, FI reduces to:

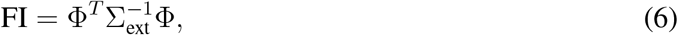

where Σ_ext_ is the input covariance matrix and 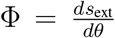 is the input gain, that is, the derivative of the stimulus input with respect to the tuning parameter *θ*. Since equation 6 is simply the *N* - dimensional analogue of equation 5 and similarly depends only on *P*_ext_, this result explains the invariance of FI to modulation for any network (as in Figure 1d).

We now seek an expression for FI in terms of only the *E* population consistent with a readout restricted to projection neurons. For the two-unit *E/I* network, the information read out from the *E* population alone is defined as [23]:

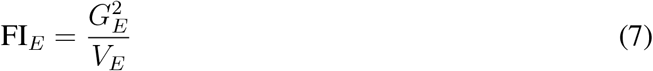

where 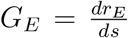 is the gain and *V*_*E*_ is the variance of the *E* population. This expression is the information analogue to a projection of the readout onto the *r*_*E*_ axis. Because it involves restricting readout to a subset of the population we refer to this as *subpopulation coding*. After some algebra we can write FI_*E*_ as

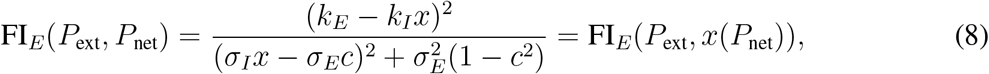

where 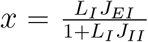. It is now apparent that the information in this subpopulation *does* depend on network state through the variable *x*, which is a function of the network parameters *P*_net_. Therefore FI_*E*_ can change with modulation (compare equations 8 and 5). Surprisingly, equation 8 reveals that this dependence on network state comes only from *L*_*I*_, *J*_*EI*_ and *J*_*II*_; that is, the information gleaned from the *E* population depends only on inputs to the network, together with the linearization of the *I* population and the *I* connectivity (Figure 3a, blue), but not explicitly on either the *E* gain or excitatory recurrent connections (Figure 3a, gray). Said differently, it is only those units which project *into* the readout population dictate the extent to which information readout changes with network state.

**Figure 3:**
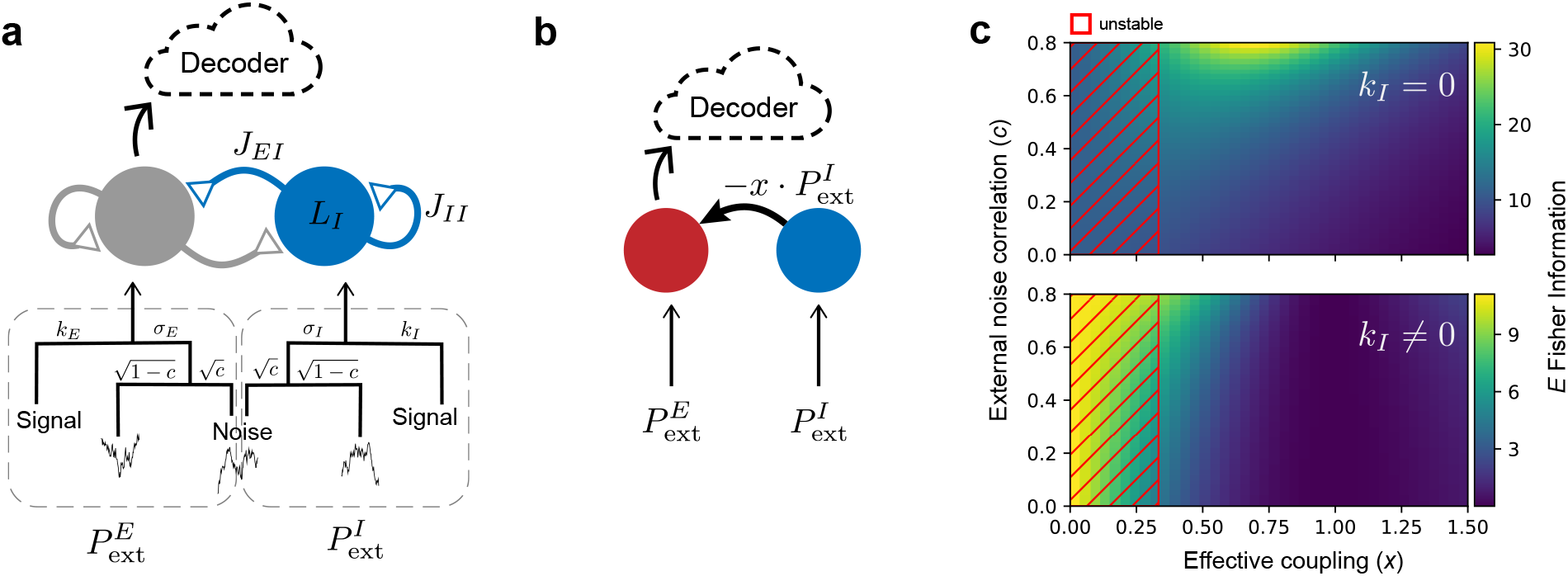
Changes in FI_*E*_ depend only on inputs to *E*. **a**. Illustration of the network parameters that affect FI_*E*_ in the network described in Figure 2 (blue). Network parameters which do not affect FI_*E*_ are in gray. *P*_ext_ has been split into *E*- and *I*-specific external inputs (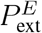 and 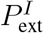,respectively). **b**. Reduced network model. FI_*E*_ depends only on feedforward inputs to *E*. **c**. FI_*E*_ as a function of *x*(*J*_*EI*_, *J*_*II*_, *L*_*I*_) and *c*. Red hatched region indicates unstable solution for a single choice of parameters 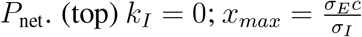. 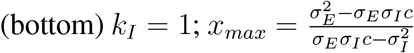. Stability line for parameters *L*_*E*_ = 10, *J*_*EE*_ = 0.18, *J*_*IE*_ = 0.24.

From the view of a linear decoder restricted to the *E* population, the recurrent network can be reduced to a feedfoward inhibition model, where *E* receives inputs *I*_0,*e*_ from external sources, and −*x*·*I*_0,*i*_ from the *I* population (Figure 3b) [30]. Here, *x* is the *effective coupling* from the inhibitory to the excitatory population (Figure 3b). Hence the effective stimulus gain of the *E* population is 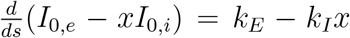 and the variance of the effective total input is 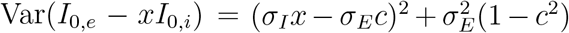, which are the numerator and denominator of FI_*E*_ (Eq. 8), respectively.

The effective coupling parameter *x* affects both the gain and the variance of the *E* population responses (Eq. 8; [30]). First suppose that the *I* population does not receive signal input, meaning that *k*_*I*_ = 0. Then FI_*E*_ can change only through the denominator, corresponding to the variance of the effective input. FI_*E*_ is maximized at 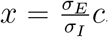, when the correlated noise from *I* population cancels the correlated component of the input noise to *E* population and minimizes the variance of the total input (Figure 3c, top). We refer to a modulation which pushes *x* closer to this value as *correlation canceling*. When there is no correlation in the inputs to *E* and *I* populations (*c* = 0), *I* merely contributes noise to *E* through *σ*_*I*_*x*. In this case, FI_*E*_ is maximal when *x* = 0, meaning that there is no projection from *I* to *E*.

By contrast, when *I* receives signal input (*k*_*I*_ *>* 0), *I* has a subtractive effect on the *E* stimulus response since *I* projects the same signal to *E* with a negative sign (Figure 3b). Therefore modulating the gain is now weighted against affecting variability to enhance information. For *x* small, reducing *x* further to minimize the subtractive impact on the *E* response is optimal, even if correlations are large (Figure 3c, bottom). As a result we label our network as being in the *gain reduction* regime. However, this brings up an important constraint since network inhibition plays an important stabilizing role as well. Thus, any changes in *x* must ensure network stability as well (Figure 3c, hatched region). Note that while we have reduced the information dependence to a single network hyper-parameter, *x*, the network’s stability depends on *all* of the network parameters *P*_net_.

The above analysis of a two-unit network corresponds to a homogeneous neuron population with identical tuning dependence on a stimulus variable. Next, we consider a population of neurons with distributed tuning preference over the encoded range of a stimulus variable, such as orientation.

### Subpopulation codes with distributed tuning

We return to a recurrent network in which *N* excitatory units connect to *N* inhibitory units (as in Figure 1a). A stimulus such as an oriented visual grating is now given by *s*_ext_(*θ*) with each unit’s preferred tuning parameter corresponding to a particular orientation *θ*.

We again consider FI_*E*_ as in the previous section, a decoder observing only the *E* population, which for a collection of *N* excitatory units takes the form (Methods)

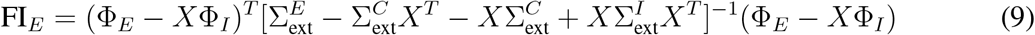

where Φ_*α*_ is the input gain Φ restricted to population 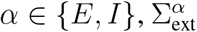 is the input covariance matrix to *α* and 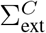 denotes the input covariance between *E* and *I*, and 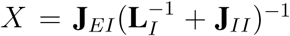 is the effective coupling matrix from the inhibitory to the excitatory population (Figure 3b). Equation 9 is simply the *N* -dimensional analogue of equation 8, thereby confirming that our preceding analysis extends to arbitrary dimension in FI_*E*_ [11, 12]. What again distinguishes FI from FI_*E*_ is that the former depends only on the structure of the input statistics to the network whereas the latter depends additionally on those network parameters restricted to the *I* population, **L**_*I*_, **J**_*EI*_ and **J**_*II*_. In particular, if we look back at the ring network which motivated this study (Figure 1), FI_*E*_ now increases.

By expanding the dimension of our recurrent network we have expanded the space of possible moduli. A complete characterization of the whole parameter space is out of the scope of this work, we are interested here in whether the mechanisms for increasing FI_*E*_ observed in the two-unit *E/I* network relate to phenomena observable in the *N* -dimensional network, namely gain modulation and correlation cancellation. We model a neuromodulatory effect as a transient rescaling in synaptic weights, **J**_*EI*_ [31] (Figure 4ai,aii; blue connections). Reducing **J**_*EI*_ here diminishes the effective projection from *I* to *E* thereby disinhibiting *E*.

**Figure 4:**
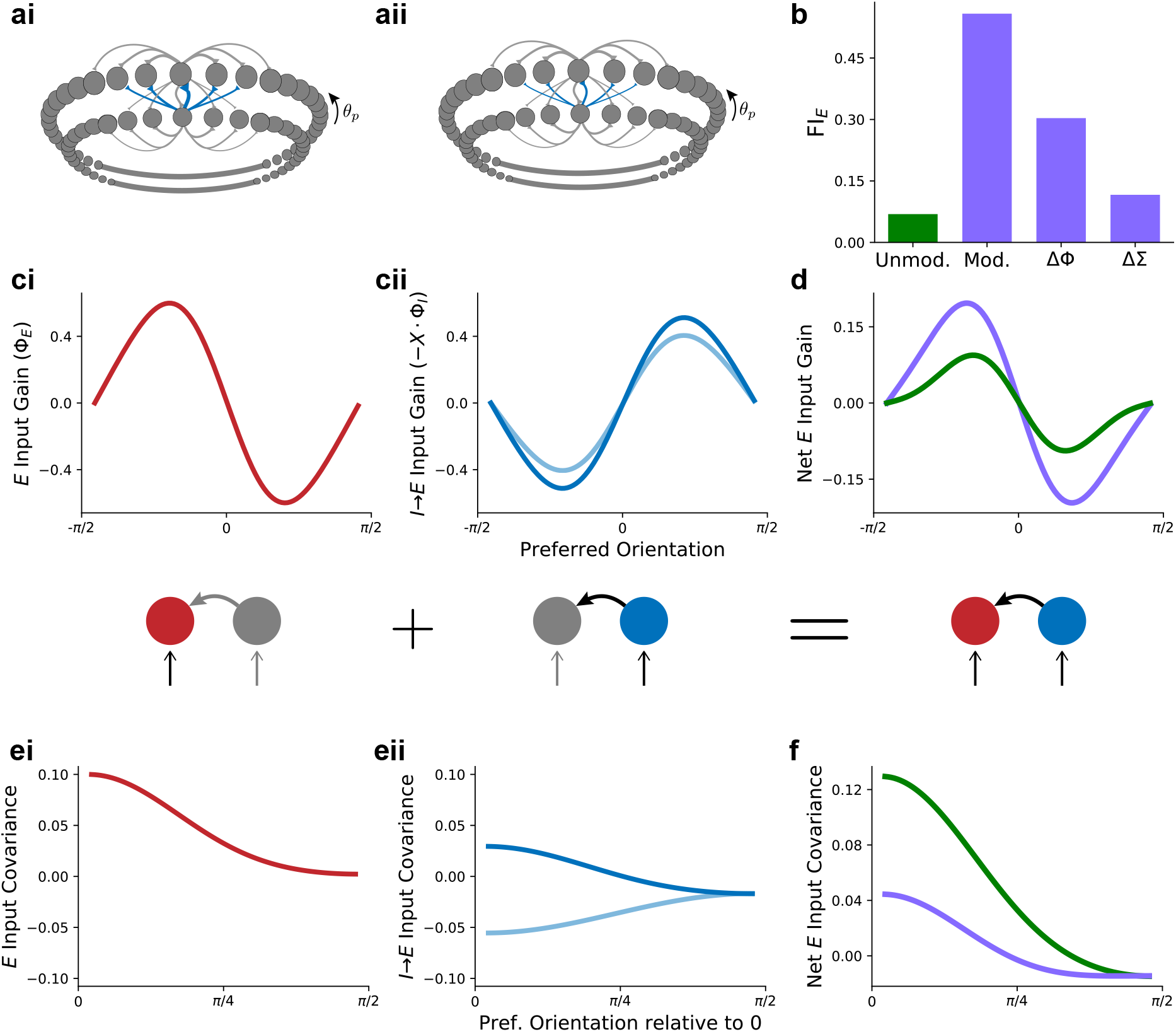
Modulation affects inputs to *E*. **a**. Network schematic of modulation via changes in synaptic strength. **b**. FI_*E*_ for unmodulated (green), modulated (mod.), modulated with changes in gain only (ΔΦ) and modulated with changes in covariance only (ΔΣ). **c-f**. 0^*o*^ is chosen to be the unit whose preferred orientation matches the peak of the stimulus. **c**. (i) External input gain *to* E units is constant with modulation. (ii) *I* inputs to *E* are scaled by *X* and depend on modulation (light blue). Schematics as in Figure 3b indicate which network elements determine the plotted values above and below. **d**. Net external input gain to *E* before (green) and after (purple) modulation. **e** (i) Illustration of a row from the input covariance to *E*. (ii) The effective input noise from *I* to 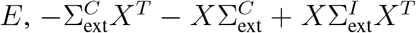, before (dark blue) and with (light blue) modulation. **f**. Total input covariance to *E*. Colors as in (d).

We again see that the net input gain and covariance to the *E* population produces FI_*E*_. To understand the impact each has on FI_*E*_ we changed *X* in equation 9 from unmodulated (*X*^*U*^) to modulated (*X*^*M*^) in either the net input gain (Φ_*E*_ − *X*Φ_*I*_) or the covariance term 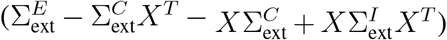 alone while keeping the other *X*’s fixed in the unmodulated state. Both substitutions result in increased FI_*E*_ (Figure 4b), with a change in the gain term resulting in an approximate three-fold increase and a change in the covariance resulting in an approximate doubling of information. What this illustrates is that both gain and covariance changes play significant roles in improving information readout. Furthermore, their joint effect is multiplicative, as the net increase in FI_*E*_ with modulation is almost six-fold.

We now explore the mechanism by which *X* increases gain and decreases covariance. In our framework external inputs to the network do not change with a state modulation by assumption, thus the *E* input gain Φ_*E*_ and input covariance 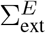 are fixed across modulation (Figure 4ci,ei). Since **J**_*EI*_ decreases in magnitude, the suppression of signal gain from *I* inputs (*X*Φ_*I*_) is reduced (Figure 4cii) leading to an increase in the net *E* input gain (Figure 4d). Additionally, this modulation resulted in a reduction of the projected *I* covariance, that is, the variability in *E* due to the *I* to *E* connection (Figure 4eii). In particular, the modulation of the projected *I* covariance results in the partial cancellation of correlations and a net reduction in *E* variability (Figure 4f). These joint improvements in signal and noise therefore combined nonlinearly to induce the increase in FI_*E*_ (Figure 4b). In conclusion, the two-unit model accurately anticipated how modulation can affect population codes through a combined effect on the effective gain and covariance.

Previous information-theoretic studies have identified a source of correlated variability, termed differential correlations, which causes information to saturate in a population [37, 24]. The differential correlation imposes an upper bound on the efficacy of population codes. As we show in the Supplemental, differential correlations limit FI_*E*_ as well. However, modulation can still increase information in a neural population provided the system has not yet saturated the bound (Figure S1). Recent studies have attempted to estimate information-limiting correlations in primary visual cortex in mouse and monkey, arguing for the presence of differential correlations which bound the information encodable by the neural population [36, 22]. However a powerful recent study recording from tens of thousands of neurons across thousands of trials did not find evidence for information saturation in mouse V1 [51]. All of these studies have considered the collective activity of V1 as the full population, encoding an oriented visual grating, say, and relating it to behavioral performance. Previous anatomical experiments have shown that feedforward projections from V1 are rather segmented into partially overlapping patches [46]. In our view, a decoder would represent the downstream area connected to a particular upstream patch. This partitioning of cortical areas could explain how, even in the presence of information-limiting correlations, neural processing can be kept away from the information saturation bound and therefore modulated with network state.

### The implications of subpopulation codes for divergent cortical pathways

The results we have described thus far are not unique to a partitioning of *E/I* networks by excitatory and inhibitory neurons. Rather, they are defined in terms of readout (or: observed) and non-readout (unobserved) populations. Given the extensive branching of corticocortical projections, this differentiation is relevant for pathways diverging from within the same cortical area to project to disjoint downstream targets. In this way, local recurrent connections or parallel yet interconnected pathways can influence one another’s information processing.

The benefits of our framework in analyzing information flow through the circuit can be seen by considering two recurrently connected *E* populations stabilized by a single *I* population (Figure 5ai,bi). A decoder reading out from both *E* populations is equivalent to FI_*E*_ computed in the preceding sections. As before, FI_*E*_ is invariant to changes in *E* activity and to all *E* connections. This can easily be seen from equation 9 applied to this network in which 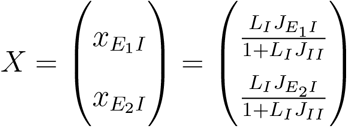. What this formulation nicely reveals is that the components of *X* are simply the two *x*’s for the subnetworks *E*_1_*/I* and *E*_2_*/I* (see effective connectivity diagrams, Figure 5aii axes). In particular, each component of *X* is the effective projection from *I* into each element *E*_1_, *E*_2_ of the readout population. Similar to the case of only one *E* population (Figure 3), if we examine the information landscape with the chosen parameter set, reducing 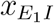 and 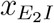 would lead to an increase in information, consistent with a disinhibitory mechanism (Figure 5aii). For example, movement toward the 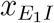 axis would be achieved by weakening the *I* → *E*_2_ connection (the alternative of changing *L*_*I*_ or *J*_*II*_ would of course affect both 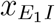 and 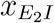). However it must be ensured that this modulation does not lead to instabilities in the network, a condition which again depends on all network parameters *P*_net_ (red region, Figure 5aii).

**Figure 5:**
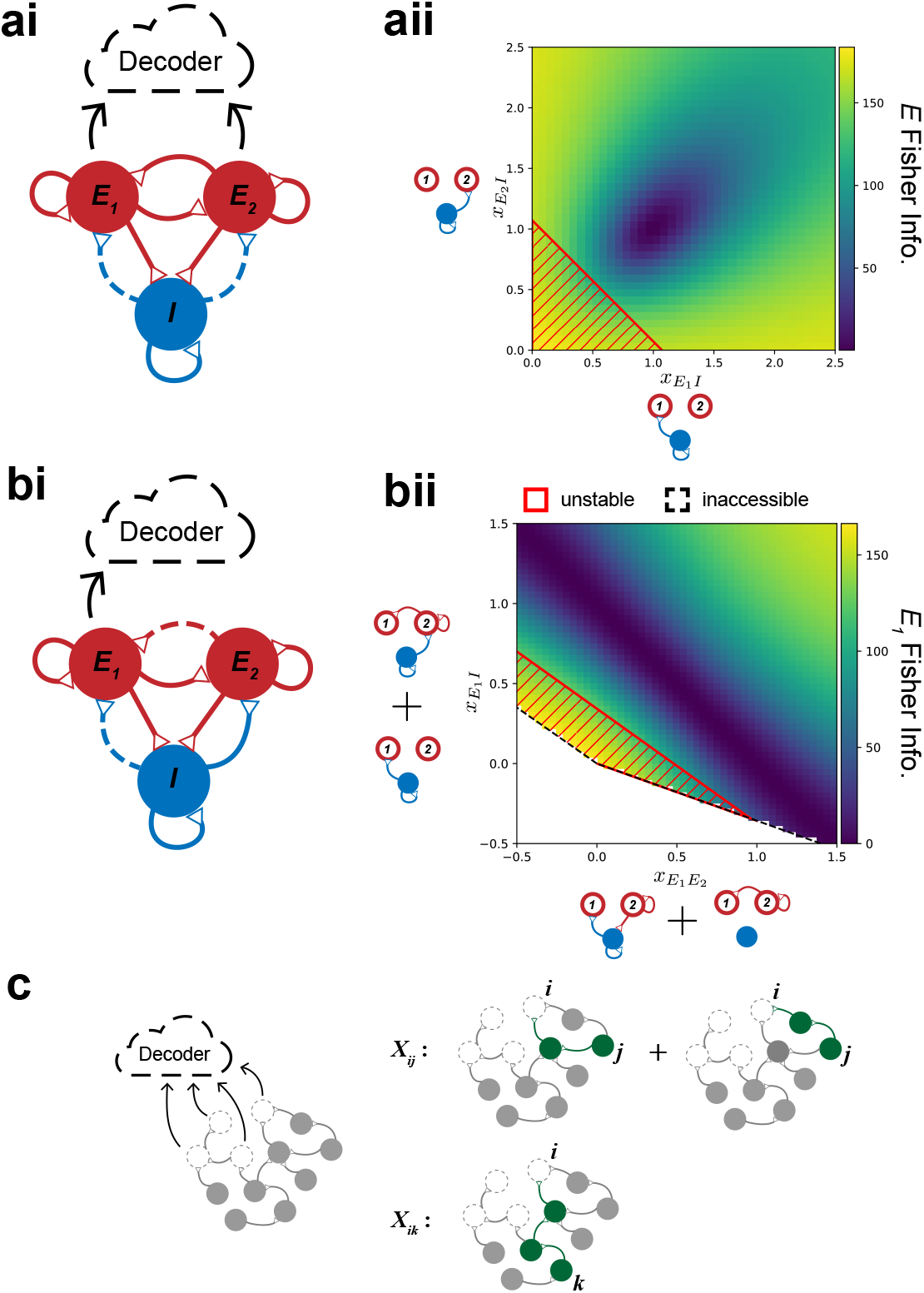
**a**. (i) Network schematic. Dashed lines illustrate which connections were varied in computing stability bounds in (ii). Readout is from both *E*_1_ and *E*_2_. (ii) FI_*E*_ for 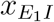 and 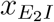 for fixed input covariance and stimulus. Axes show the effective connections which comprise each *x*. Red region indicates instability. **b**. (i) As in (ai) for readout from only *E*_1_. (ii) As in (aii) where the white region indicates inaccessible values of 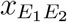 and 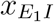 for the chosen parameter set. **c**. Illustration of the general form of elements of *X*. A generic network is shown (grey) with only a subset of units available to the decoder (grey dashed circles). Two elements of *X* are illustrated in terms of the pathways through the network which contribute to them (green).

Now suppose *E*_1_ and *E*_2_ project to different targets, and consider the downstream decoder reading out from only *E*_1_ (Figure 5bi). In this case we want to analyze 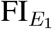 and 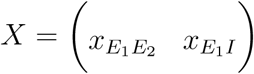 now becomes effective couplings from *E*_2_ and *I*. The two components of *X*, 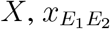 and 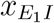, are composed of all effective paths from populations *E*_2_ and *I*, respectively, *to E*_1_ (Figure 5bii axis diagrams; Eqs. 20, 21). In this case, 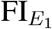 is high when 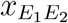 and 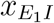 are both large or small (Figure 5bii). Therefore, information of the *E*_1_ population can be increased by jointly increasing or decreasing 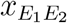 and 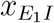, which can be achieved by, for example, strengthening or weakening 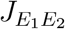 and 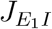 (Figure S2). Note that there is a region of inaccessible values of 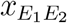 and 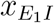 (Figure 5bii, white region), due to the restrictions Dale’s law places on the signs of the connection weights (i.e. *J*_*αE*_ ≥ 0; see Supplemental Material). Similarly, the parameter space for FI_*E*_ is restricted to positive values of 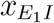 and 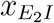 (Figure 5aii). The inaccessible region of *x*s depends on connection strengths and cellular gains (Figure 6).

**Figure 6:**
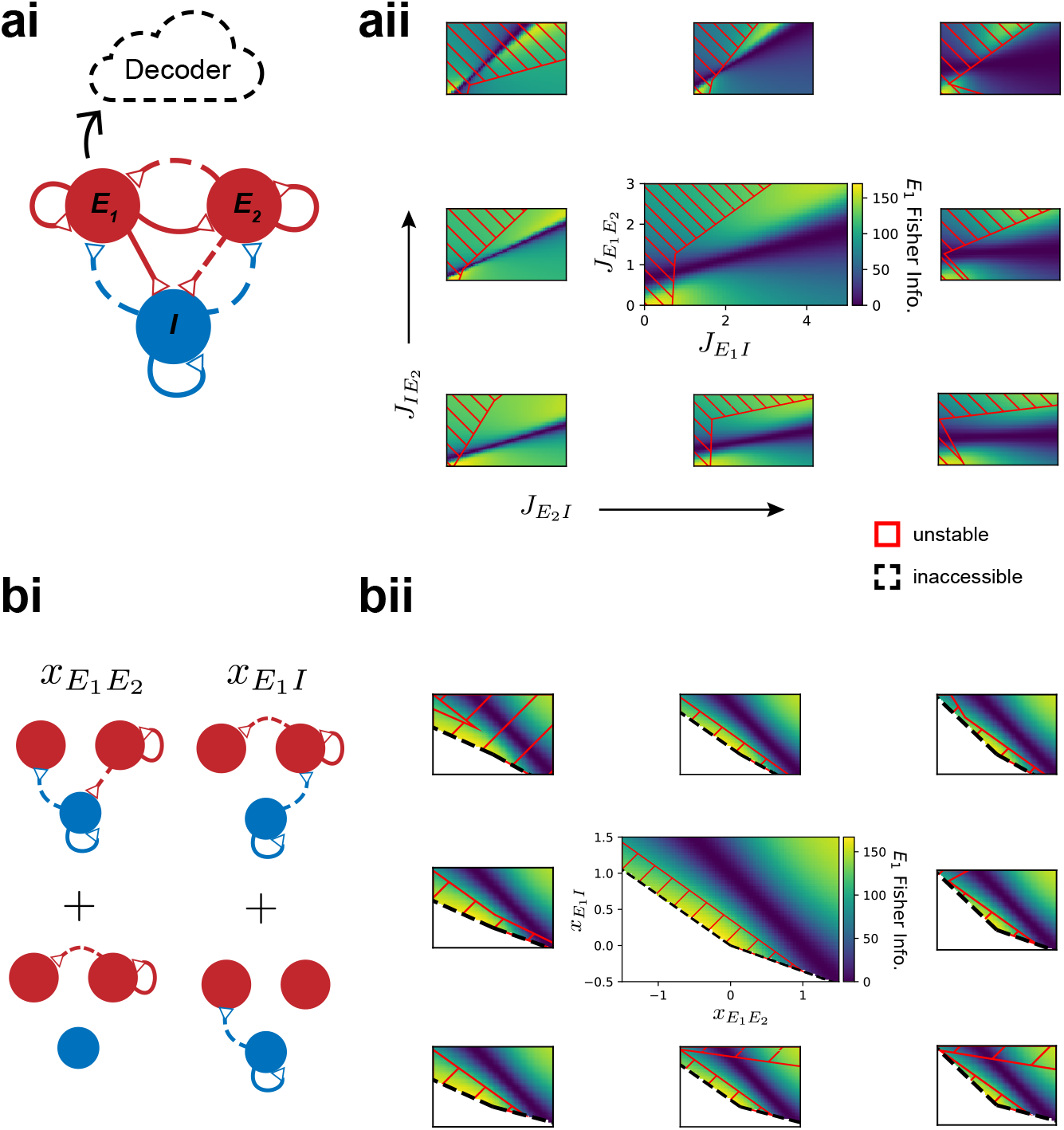
Parametric benefits of the theory. **a**. (i) Schematic of the network. Dashed lines represent connections varied within and across plots (stability boundaries computed as in Figure 5). (ii) FI_*E*_ as a function of varying connectivities (*J* ’s) for a semi-naive surf of *J* -space. Red regions indicate instability. Direct connections from *E*_2_ and *I* to *E*_1_ were varied within a plot. Connections between *E*_2_ and *I* were varied across plots. **b**. (i) Illustration of connections contributing to 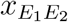 and 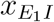 with varied connections dashed. (ii) Same data and arrangement as in (aii) plotted as a function of 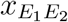 and 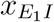. Central plot in (aii) and (bii) is same parameter set as Figure 5bii.

In a general network of multiple units, we find that *X* comprises all paths through the unobserved units together with all projections to each readout unit (Figure 5c and Supplemental Material). The analysis is similar to previous works which decompose the network response covariance into structural motifs [56, 48].

In sum, the modulation of information flow in cortical networks is affected by all input connections to the readout population. This poses a thornier issue for populations projecting to divergent targets than for *E* networks with the same decoder since activity along one pathway can influence information flow along another connected path. As we speculate in the Discussion, this result could motivate compartmentalization, or clustering, of activity in cortex.

### Parametric considerations in the theory of subpopulation codes

While we have thus far argued for the biological importance of our theory, it has a mathematical benefit as well. In order to understand the effect of modulation on information flow in a network, traditional analysis would require knowledge of all network parameters. For a network of *N* units this is *N* (*N* + 1) different parameters if we consider all possible connections (*N* ^2^) and each unit’s stimulus-response linearization (*N*). In our theory, however, if readout is restricted to *m* of the *N* total units one must only understand how modulation affects *m*(*N* − *m*) parameters, the number of elements in *X*. This can offer a significant advantage. For instance, in the preceding example we reduced the relevant parameter space from 12 to 2 when decoding from *E*_1_.

Other considerations such as system stability will yet demand full knowledge of the system. In spite of this, our theory still affords some benefit (Figure 6). We return to the question of decoding from *E*_1_ in the same network as in the preceding section (Figure 5) by varying four connection strengths (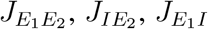 and 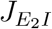; Figure 6ai). If we look at how 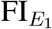 changes when viewed as a function of these four connection strengths, both the stability boundaries and the information landscapes change across panels (Figure 6aii). Describing a consistent theory for how modulation can enhance information flow in this context would be exceedingly difficult; there are still five unexplored connections. By contrast the same parameter changes replotted in terms of 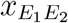 and 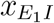 show a different picture (Figure 6bi): the information space is invariant, and only stability and accessibility boundaries change (Figure 6bii). Given that a modulation can drive a change either within or across plots, this representation affords a much clearer picture of how to modulate the network once network stability is taken into account. We comment, however, that the connection parameters we manipulated here still offer a peek behind the curtain; without knowledge of which connectivity parameters most influence *X* it would not have necessarily been *a priori* obvious to consider 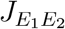 vs. 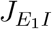 and one might have had to explore all nine connectivity values to arrive at the same conclusion.

## Discussion

In this study we explored how information within a neural population changes as a function of cortical state. We have shown that for cortical state to affect information flow in a neural circuit, a linear decoder must only observe a subset of the complete neural population, consistent with cortical anatomy in which only a subset of neurons project to downstream targets [15]. Moreover, we observed the counter-intuitive result that under changes in cortical state, it is only those neurons which are not decoded from (i.e. do not project to a downstream region) that shape linear readout from a neural population that does project downstream. This suggests a strong role for inhibition in shaping information flow in cortex, a view which is gaining broad support [18]. Furthermore, we showed that subpopulation linear Fisher information depends only on the structure of the external (e.g. thalamocortical) inputs to the network, together with the non-readout connections and input/output linearization, effectively reducing subpopulation codes to a feedforward circuit.

In the ring model with distributed tuning, we assumed that any current modulation is independent of the tuning preference of neurons (Figure 1). While tuning-dependent modulation has been used to model selective attention [8, 28], it trivially adds information to the network. Specifically, denoting current modulation as *M* (*θ*), we would have 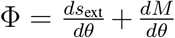. Hence, in this work we only considered modulations that do not trivially introduce information to the network, such as transient synaptic weight changes and tuning-independent current modulation.

Throughout this study we largely focused on a classic *E/I* dichotomy in which excitatory neurons conveys information to downstream regions while inhibitory neurons solely act locally. However our analysis proved relevant to other common circuit motifs such as those in which local excitatory cells shape projection cell output or divergent projection cells mutually interact. As an example, intracortical pyramidal neurons in motor cortex innervate corticostriatal cells which project from motor cortex to various subcortical and peripheral targets [50]. Additionally, deep layers of cortex are also the source of reciprocal connections between divergent projection neurons: corticothalamic neurons which project subcortically and intratelencephalic neurons which project within cortex [16]. Given the high divergence of cortical projection pathways and the preponderance of clustered network architectures across cortex, our results support an information-processing benefit to a clustered organization [29]. By limiting the interaction between neural clusters performing distinct computations, the brain could achieve better control over the degree to which activity in one cluster affects the information flow in another.

Previous studies have considered readout from only the excitatory neurons [30, 38, 23]. However these studies either did not explore the impact of modulation on the decoder [30] or considered only homogeneous tuning inputs [23], leaving open the question of whether and how these results extended to networks with distributed tuning. A recent study [38] explored the coding capacity of inhibitory neurons, thus comparing *E* vs. *I* information, and examined how changes in *E/I* connectivity (*J*_*EI*_, *J*_*IE*_) affected information content in *E* and *I* populations separately. Our results clarify why varying these connections was the only way to affect information in their network: their use of a linear system fixed the cellular gain (*L* in our theory), and the only other terms the effective connectivity (*X*) depend on are the *J* ’s.

The effects of state modulation on neural circuits are highly diverse, capable of adjusting a range of properties from cell-intrinsic - such as excitability and neurotransmitter release [14, 31] - to population-wide, such as oscillatory activity and noise correlations [54]. The mechanisms underlying state changes are equally diverse, involving many classes of neuromodulator and different sources of feedback drive [17]. Neuromodulators can be broadly distributed, such as cholinergic projections from midbrain to cortex, or highly specific, like dopaminergic targeting of individual cortical layers [53, 54]. This seemingly endless flexibility poses a challenge for determining how these moduli relate to neural processing. Our theory goes some way towards identifying the circuit components which are actually affecting neural processing. For example, we found that reducing the synaptic strength between *I* and *E* can increase gain and decrease covariance in excitatory units. These effects are mirrored by acetylcholine (ACh), which has been shown to reduce synaptic efficacy of intracortical connections [54].

Neuromodulation can affect the responsiveness of neurons in many ways that our model does not capture, for example, by reducing burst spiking and altering firing adaptation [17]. Of course, the omission of a spiking mechanism is a clear limitation of rate models, and future work should address the way in which modulation of spiking properties affects neural coding across states. Our choice of firing rate models are nevertheless able to capture the main effects observed in many experimental paradigms such as primate electrophysiology experiments in which analysis is performed on spike counts [5], or calcium imaging experiments [52]. In fact, we anticipate that the spatially broad recording capacity of modern calcium imaging, together with neural subtype indicators will provide the data our model has identified as important to understanding the circuit mechanisms of information modulation, namely, the activity of local (largely inhibitory) interneurons.

Our results also highlight the general difficulty in assigning a functional role to specific network components in affecting information flow. If we consider an arbitrary connection *J*_*EI*_ this can emerge in multiple paths from the non-readout to the readout population, thereby affecting multiple values in *X* (Supplemental Material; Figure S2). Similarly, the linearization *L*_*α*_ of unit *α* contributes to all values of *X* in which connections from unit *α* are present. These issues are natural consequences of the recurrent nature of cortical circuits. In spite of this, the ability to map a recurrent circuit to an effectively feedforward model with an interpretable hyperparameter, *X*, will facilitate uncovering the mechanics of state-dependent information modulation.

Understanding how neural processing depends on brain state is a major goal in systems neuroscience [7]. Even in simple subcortical systems like the crab stomatogastric ganglion in which the full connectivity structure is known, a modulus’s effect on circuit dynamics is still difficult to generalize as it depends upon the relationship between the network state and the nature of the modulus [32]. Expanding to cortical circuits of much larger size poses a daunting challenge, however, our theory significantly reduces the space of parameters which need to be measured, as well as provides a degree of interpretability by representing information flow in terms of effective pathways within a network. Our results highlight the importance of locally projecting neurons in shaping the information in neurons that project to downstream areas.

## Acknowledgements

The authors thank Jeff Dunworth for assistance on an early version of this project, Hannah Bos and Ross Williamson for helpful discussions, Olivia Gozel and Gregory Handy for a close-read of the manuscript, and Tom Bonamici for graphics advice.

## Methods

Code will be made available on the author’s github upon publication: github.com/mpgetz

### Linear theory

We considered a linearization of a nonlinear firing rate model (equation 2). For sufficiently small noise, equation 2 can be linearized around the steady state rate, with fluctuations in rate around the operating point given by an *N* -dimensional extension of traditional linear response theory:

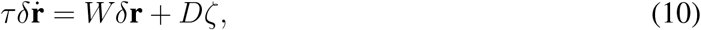

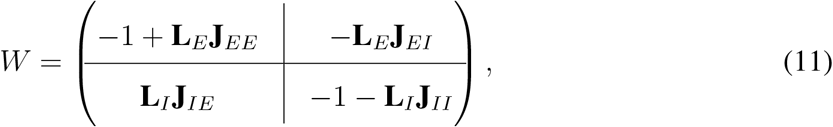

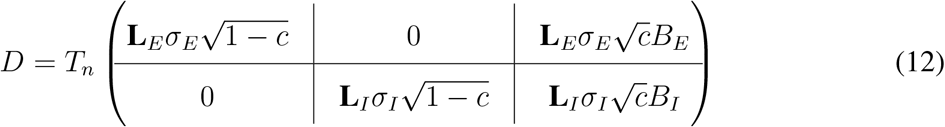

where *ζ* is the (*N* + *ν*)-dimensional vector of external input noise (here *ν* = 1 is the dimension of the shared external input variability), *W* is an *N* ×*N* matrix of effective weights, *D* is *N* ×(*N* +*ν*) matrix which scales the noise terms, *B*_*α*_ is the *N*_*α*_×*ν* matrix which determines the input covariance structure, *σ*_*α*_ is the amplitude of the noise to a given unit, 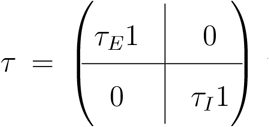 where 1 is the *N*_*α*_ × *N*_*α*_ identity matrix. **L**_*α*_ is the diagonal matrix of derivatives of the transfer function at the steady state rate whose *i*^*th*^ diagonal element is given by

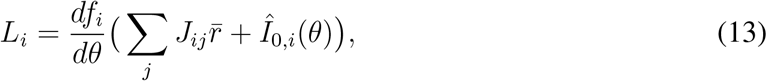

with 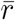 the steady-state solution to equation 2 and *Î*_0,*i*_(*θ*) = *b* + *s*_ext_(*θ*). In order to avoid injecting pure white noise into the system, we consider a temporally smoothed noise process

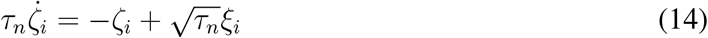

where *ξ*_*n*_ is a white noise process and *i* is an index over all possible noise sources. Consequently, *T*_*n*_ is a time-scaling constant given by 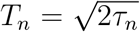. This linearized stochastic system then enables us to estimate the full covariance matrix of the spatially extended model [10]. In particular, the covariance matrix is given by Σ = *W* ^*−*1^*D*(*W* ^*−*1^*D*)^*T*^. It should be noted we have written the connectivity parameters **J** as positive values; the negative sign of inhibition is made explicit in the dynamical equations and carried through the relevant derivations.

### Gain calclulation

The population response gain is given in general by the derivative of a population’s response with respect to an input parameter. In the reduced *E/I* model we have *s*_ext,*α*_ = *k*_*α*_*s* such that 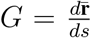. In the ring network, the parameter of interest is the angular variable *θ* such that 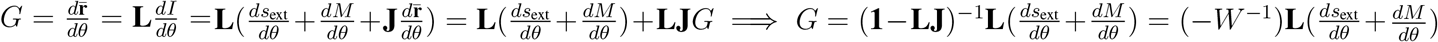. Here we have let *I* equal the argument of *f* in equation 2. We have generalized this derivation with the inclusion of a current modulus *M* (*θ*) such that *I* = **J** · **r** + *I*_0_ + *M* (*θ*) where *I*_0_ is given by equation 3; a transient weight change would simply affect **J**. Finally, letting 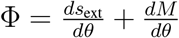 we arrive at the following compact expression: *G* = −*W* ^*−*1^**L**Φ.

### Fisher information analysis

#### Full FI

As mentioned above we write the (long time) covariance matrix Σ = (*W* ^*−*1^*D*)(*W* ^*−*1^*D*)^*T*^ = *W* ^*−*1^*DD*^*T*^ (*W* ^*−*1^)^*T*^. Notice that we can rewrite *D* such that *D* = **L***D*_ext_ where 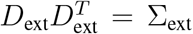. Hence, we have

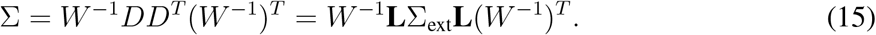

The linear Fisher information (FI) is thus given by

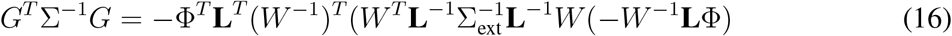

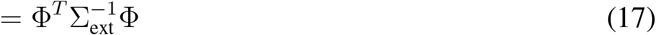

since **L** = **L**^*T*^.

### FI_*E*_ derivation

We computed FI for the *E* population similarly by calculating 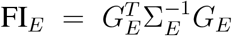 as follows. Partitioning the input covariance matrix into a block structure such that 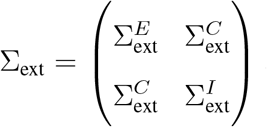 and applying equation 15 leads to

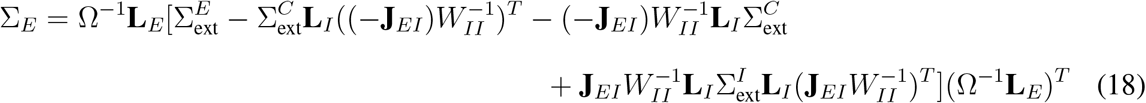

where 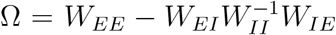 and we have made use of the fact that *W*_*EI*_ = −**L**_*E*_**J**_*EI*_. By a similar argument we have that

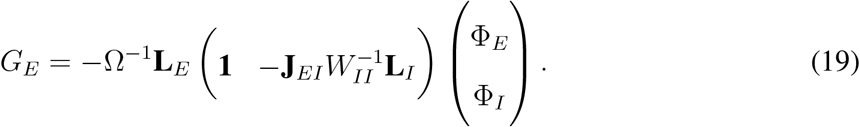

Plugging these two expressions into the FI_*E*_ equation at the beginning of this section and identifying the term 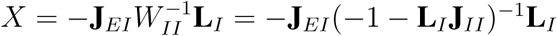 gives the final equation 9.

### Subpopulation codes in general: FI_*α*_

The derivation shown above in the section “FI_*E*_ derivation” is flexible to shifts in the block structure. Thus for any subpopulation *α* of the full network the same arguments apply for a partitioning of the inputs with the mappings *E* → *α* and *I* → *U* where the unobserved network elements are denoted by *U*.

### Derivation of *X* for divergent *E* populations

Here we derive *X* for FI_*E*_ and 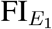 in the two recurrently connected *E* populations and single *I* population (Figure 5). From equation 9 we have that, for FI_*E*_, 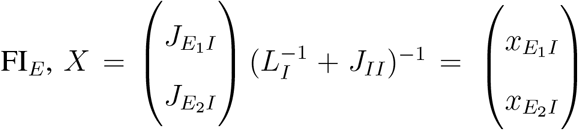 where, since *L*_*I*_ and *J*_*II*_ are scalars, 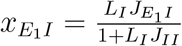 and 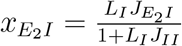.

We now derive 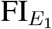 after the preceding section. Let *U* = {*E*_2_, *I*}. Then for *E*_1_ readout, 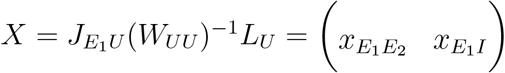 where

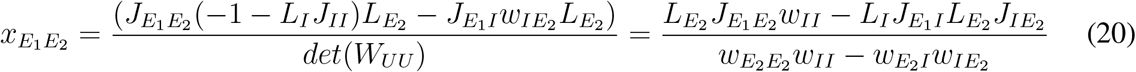

and

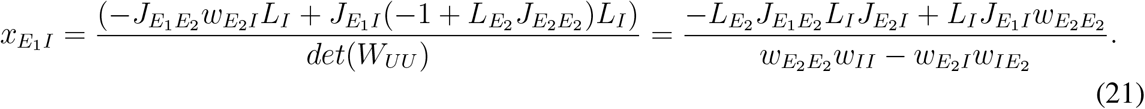

Here we have written *w*_*αβ*_ to denote the individual elements of the *W* matrix for this network, defined in equation 11 (for analysis of this network see: Supplemental Material). In particular, *w*_*αβ*_ = −*δ*_*αβ*_ + *L*_*α*_*J*_*αβ*_ where *δ* is the Dirac delta function.

### Model parameters

#### *E/I* network (Figure 2)

The response distributions were given by *r* = −*W* ^*−*1^**L***I* and Σ = (*W* ^*−*1^**L**)Σ_ext_(*W* ^*−*1^**L**)^*T*^. Here we used the *W* notation defined along with equation 11. Letting *A* = −*W* ^*−*1^**L** with superscripts to denote the relevant figure panel in which the parameter set was used, we have: 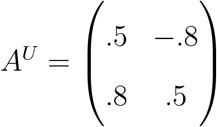, 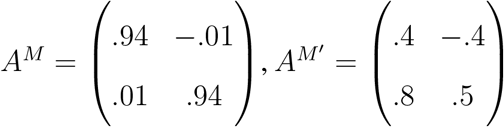. Input drive 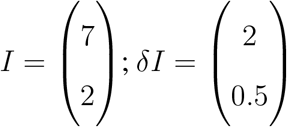; input covariance 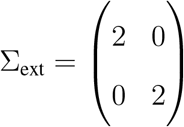. Additionally, *L*’s and *J* ’s which also solve the equations are given in Table 1.

**Table 1.**
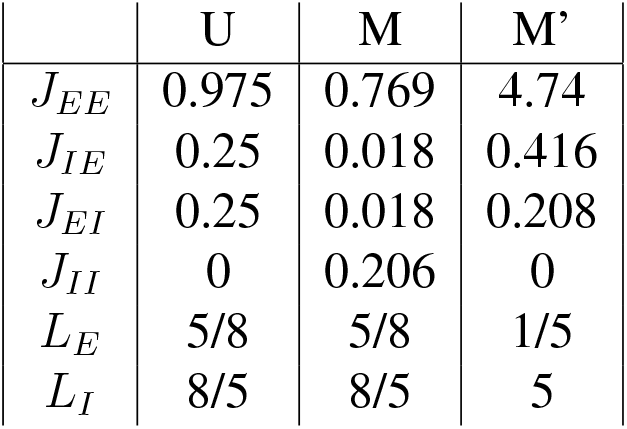

|  | U | M | M' |
| --- | --- | --- | --- |
| $J_{EE}$ | 0.975 | 0.769 | 4.74 |
| $J_{IE}$ | 0.25 | 0.018 | 0.416 |
| $J_{EI}$ | 0.25 | 0.018 | 0.208 |
| $J_{II}$ | 0 | 0.206 | 0 |
| $L_E$ | 5/8 | 5/8 | 1/5 |
| $L_I$ | 8/5 | 8/5 | 5 |

### Ring network (Figures 1 and 4)

The ring model consisted of *N*_*E*_ = 180 *E* units and *N*_*I*_ = 180 *I* units where location on the ring corresponded to a unit’s preferred tuning *θ*. Parameters were derived in part from Rubin *et al*. [44].

For all units the transfer function is threshold quadratic: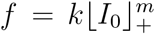. Here *k* = 0.04 and *m* = 2. The stimulus is a wrapped gaussian with amplitude *c*_*α*_: *s*_ext_(*θ*) = *c*_*α*_*g*(*θ*; *θ*_0_, *σ*_ext_) where

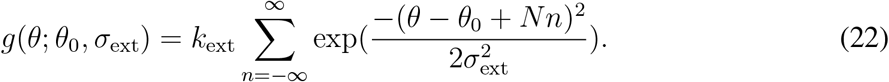

The baseline input *b* = 10, *k*_ext_ = 40 and the stimulus was centered at *θ*_0_ = 90^*o*^. Modulation was introduced as a uniform current bias *M* (*θ*) = *µ* so that *I* = *I*_0_ + *µ*_*α*_ with *µ*_*E*_ = −1, *µ*_*I*_ = 2 (Figure 1), *µ*_*E*_ = −2, *µ*_*I*_ = 4 (Figure S1). Modulation in Figure 4 was modeled as a change in connectivity weight (see Table 2).

**Table 2:** Superscript *M* in the Figure value denotes the value to which a parameter changed with modulation.

|  | Value | Units | Figure |
| --- | --- | --- | --- |
| $\tau_E$ | 20 | msec | 1, 4 |
| $\tau_I$ | 10 | msec | 1, 4 |
| $\tau_n$ | 1 | msec | 1, 4 |
| $J_{EI}$ | 0.023 | | 1 |
| | 0.03 | | $4^M$ |
|  | 0.043 |  | 4 |
| $J_{EE}$ | 0.044 | | 1 |
|  | 0.027 |  | 4 |
| $J_{IE}$ | 0.042 | | 1 |
|  | 0.03 |  | 4 |
| $J_{II}$ | 0.018 | | 1 |
|  | 0.042 |  | 4 |
| $\sigma_E$ | 1 | | |
| $\sigma_I$ | 1 | | |
| $\sigma_{ext}$ | $30^\circ$ | | |
| | $40^\circ$ | | 4 |
| $\sigma_E$ | $30^\circ$ | | |
| $c_E$ | 0.1 | | |
| $\sigma_I$ | $20^\circ$ | | |
| $c_I$ | 0.5 | | |
| $\sigma_C$ | $30^\circ$ | | |
| $c_C$ | 0.2 | | |

Connectivity decayed with distance, given by *g*(*θ*; *θ*_*α*_, *σ*_*αβ*_) with *σ*_*αβ*_ = 32^*o*^ for all *α, β* ∈ {*E, I*}. Note in the figures, angles were expressed in radians.

The input covariances partitioned as in equation 18 were given by: 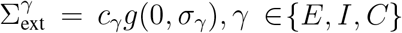 (see Table 2).

#### *E*_1_*/E*_2_ network (Figures 5, 6)

We used equation 9 to generate the plots of FI_*E*_ vs. *x*. Input variance was 0.003025 while input covariance was 0.00242. To compute the stability boundaries in Figures 5 and 6, unless varied explicitly in the Figure, weights were set to 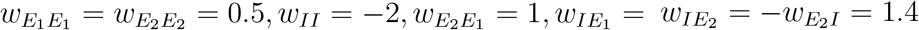 (*w*_*αβ*_ notation defined in Methods: Derivation of *X* for divergent *E* populations).

## Supplemental Material

Here we provide detail of the analyses of divergent excitatory pathways (Figure 5, 6) including stability and accessibility bounds, a discussion of differential correlations in our model [37, 24] and a brief description of the general decomposition of *X* into paths through the network (Figure 5).

### Impact of low-rank variability on modulation of subpopulation codes

Numerous experimental studies have demonstrated that the covariance structure in cortical circuits contains a significant low-rank component [20]. We therefore consider a covariance matrix which decomposes into full and low-rank components Σ = Σ_0_ + *ϵvv*^*T*^. Here Σ_0_ is full-rank and *v* is an *N* -dimensional vector. We denote the elements of *v* which act on the *E* population by the vector *v*_*E*_. Computing FI_*E*_ for this covariance structure results in (see: Supplemental, FI_*E*_ for low-rank covariance)

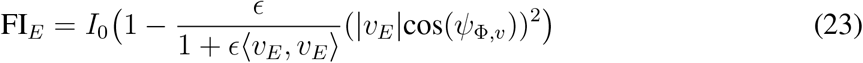

where *ψ*_Φ,*v*_ denotes the angle between Φ and *v*. We have defined 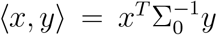 and *I*_0_ = Φ^*T*^ (Σ_0_)Φ. Thus we see two potential (nontrivial) mechanisms for enhancing information flow: through changes in *I*_0_ or by reducing cos(*ψ*_Φ,*v*_). As the numerical results explored yielded either parallel or orthogonal vectors, we focus here only on the first mechanism. In particular, let *v* be the simplest case: the unit vector (Figure S1A, top). A modulation in the form of a constant input bias to all cells in the network model described in the previous section reduces correlations for nearby units while slightly increasing correlations for dissimilarly tuned units (Figure S1B, solid lines, green to magenta). This modulation results in a linear increase in FI_*E*_ as a function of *I*_0_ (Figure S1C, solid line, green to magenta).

Equation 23 allows us to additionally consider an important case for *v*. It has been shown that the nature of correlated variability which limits information in a neural network is that which causes a shift in the population’s response in the direction of the encoded variable [37]. These are called differential correlations as for a population response **r** they take the form of 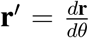 (other studies have often used the notation *f* instead of **r** [37]). To this end consider a network in the presence of differential correlations such that *v* = Φ and equation 23 simplifies to

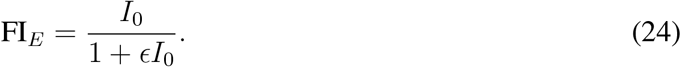

Thus, as *I*_0_ → ∞, information in subpopulation codes saturates to 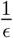 (compare curves in Figure S1C). To see how differential correlations affect modulation of subpopulation codes we introduced *v*_*α*_ = Φ_*α*_ into our model network (Figure S1A, bottom). The same modulation applied to this network resulted in an almost identical change in correlations (Figure S1B, dashed lines, green to magenta). However despite the similarities in the output correlation structures (Figure S1B), the *E*-population readout, as well as the change in the readout, differs significantly (Figure S1C, dashed line, green to magenta). Thus it is not enough to know how the firing rate statistics of a network change with modulation. Instead, one must understand the nature of the output projections of the network in question, together with its inputs, to get the full picture of how modulation is affecting information flow.

In summary, subpopulation codes can be modulated in the presence of information-limiting correlations as long as the network has not saturated the 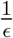 bound. Away from saturation, the results of our previous analyses for population codes hold. This is because *I*_0_ is still subject to the modulatory effects described above.

**Figure S1:**
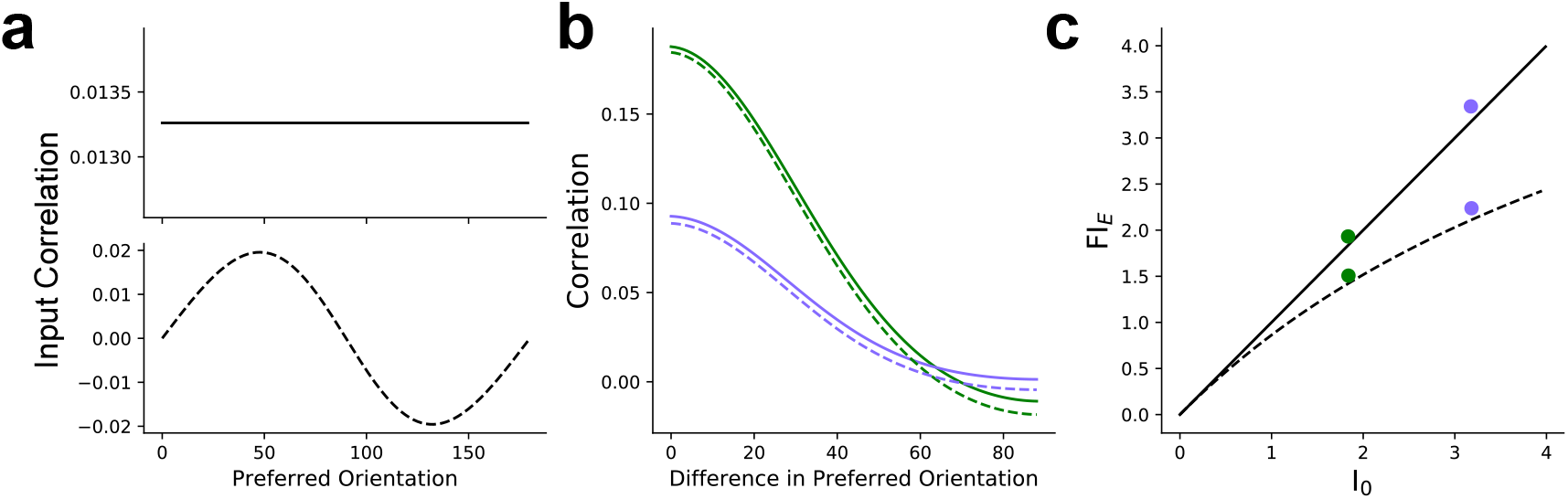
**a**. Input correlation structures. **b**. Output correlations for two different structures of input variability. Green: unmodulated; purple: modulated. **c**. Corresponding changes in FI_*E*_ for correlations in (b). Dashed line: differential correlations; solid line: non-differential correlations. *E* in equation 24 was 0.3.

### FI_*E*_ for low-rank covariance

Here we specify a low-rank component of the input covariance such that Σ_ext_ = Σ_0_ + *vv*^*T*^ and again denote *v*_*α*_ the elements of *v* which act on population *α*. Then

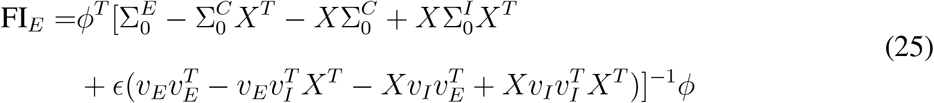

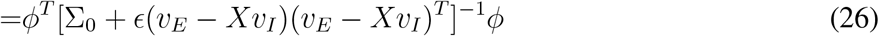

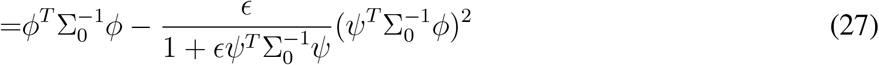

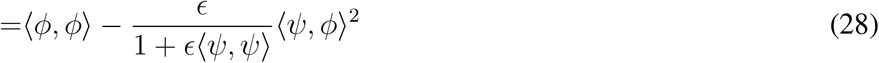

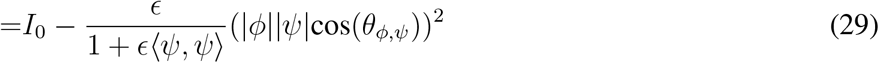

where *ψ* = *v*_*E*_ − *Xv*_*I*_ and we denoted *φ* = Φ_*E*_ − *X*Φ_*I*_. As above we let 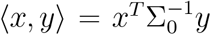 and 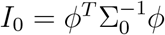.

With the substitution *v*_*α*_ = Φ_*α*_ we have in the last equation above that *ψ* = *φ* and cos(*θ*_*ϕ,ϕ*_) = 1 which results in equation 24.

**Table S1:**
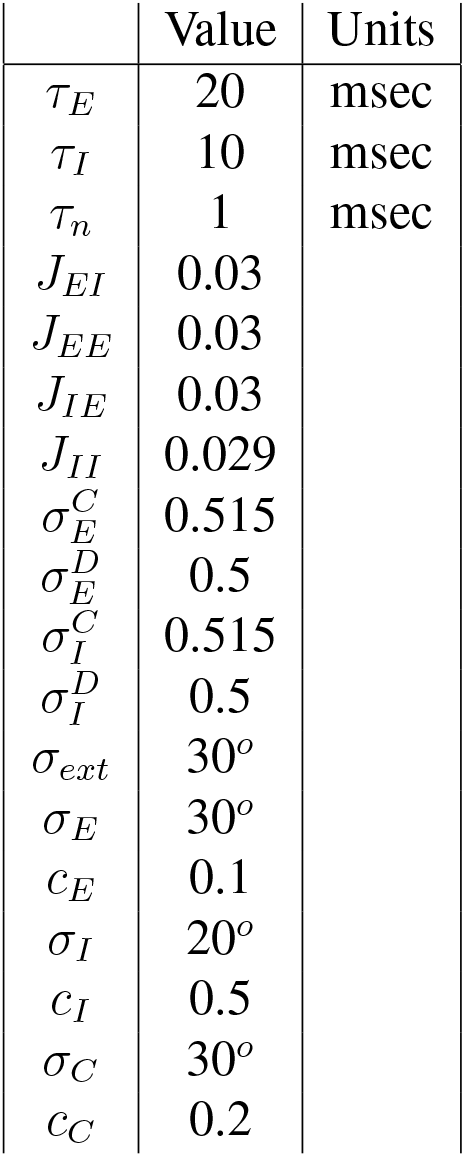
Figure S1 Parameters. Variable superscript C denotes constant modulation case, D denotes differential correlation case.

### Analysis of Divergent Excitatory Pathways

Here we expand our discussion of the *E*_1_ − *E*_2_ − *I* network described in Figure 5. In particular we outline the stability conditions analyzed and limitations on attainable values of *x*_1_ and *x*_2_. We remind the reader of the system dynamics:

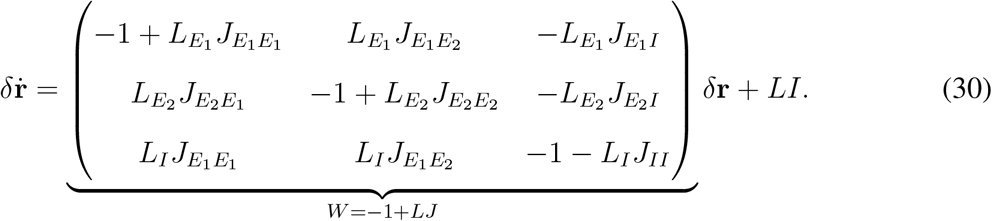

*X* for a linear decoder restricted to population *E*_1_ then has components

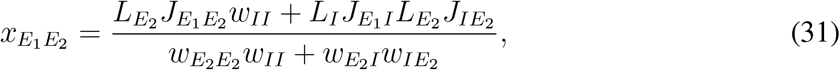

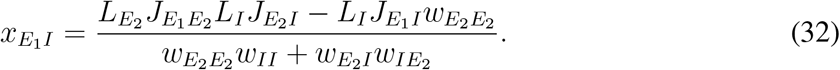

where *w*_*αβ*_ is the element of *W* given by *w*_*αβ*_ = −*δ*_*αβ*_ + *L*_*α*_*J*_*αβ*_ with *δ* the Dirac delta function and *α, β* ∈ {*E*_1_, *E*_2_, *I*}. Since we have assumed our system is in a steady-state and linearizable regime we can apply the Routh-Hurwitz stability criteria to our linearized network, which requires three conditions be satisfied (note we have not assumed a sign for any of the *w*_*αβ*_’s):

1. 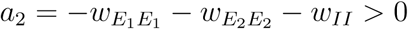
2. 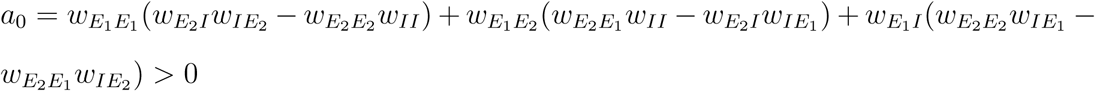
3. 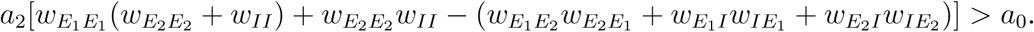.

Next we prove bounds on *x*_1_ and *x*_2_ for given parameter sets. We will use the following shorthand: 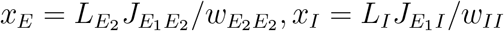 and *x*_*βα*_ = *L*_*α*_*J*_*βα*_*/w*_*αα*_, *α, β* ∈ {*E*_2_, *I*} (i.e. the *x*’s written for the corresponding monosynaptic connections). Finally, to make the notation clearer we write 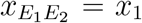 and 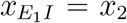. First we prove the general claim that *x*_1_, *x*_2_ cannot both be negative. More precisely, *x*_1_ *<* 0 =⇒ *x*_2_ ≥ 0 and *x*_2_ *<* 0 =⇒ *x*_1_ ≥ 0. Rearranging terms in equations 31 and 32 we can write:

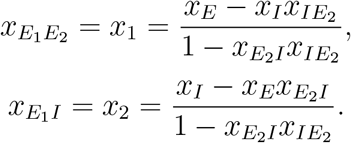

Note that *x*_*I*_ ≥ 0 since 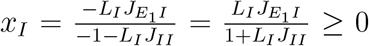, and similarly for 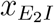. By contrast, *x*_*E*_ is unrestricted.

We will consider two cases. For readability we write 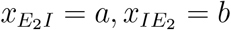. In this way

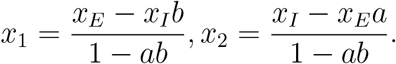

Case 1: 1 − *ab >* 0. First let *b* ≥ 0. Then

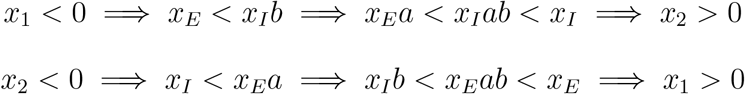

where the last inequality follows from the assumption that 1 *> ab*. Now suppose *b <* 0. But then *x*_1_ ∝ *x*_*E*_ + *x*_*I*_|*b*| *<* 0 ⇐⇒ *x*_*E*_ *<* 0 =⇒ *x*_2_ ∝ *x*_*I*_ − *x*_*E*_*a* = *x*_*I*_ + |*x*_*E*_|*a >* 0. Clearly *x*_2_ *<* 0 requires *x*_*E*_ *>* 0 so this condition is immediate.

Case 2: 1 − *ab <* 0. Note that this assumption is true iff *a, b >* 0 since *a* ≥ 0.

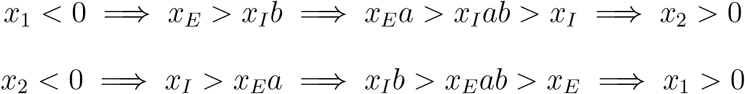

where now the last inequality follows from the assumption that 1 *< ab*.

The above arguments illustrate parameter-independent bounds on *x*_1_ and *x*_2_ for this particular system. However there are also parameter-dependent bounds on *x*_1_ and *x*_2_ (Figure 6, variability in the white regions). The main text focused on bounds due to restrictions on the connection (*J*) values; these arise from the fact that excitation and inhibition necessarily have positive and negative values, respectively. Equations 31 and 32 share the same denominator, which we denote *d*. In each panel in Figure 5 we varied only 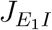 and 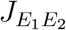 (see also Figure S2). Thus equations 31 and 32 can be written in the form:

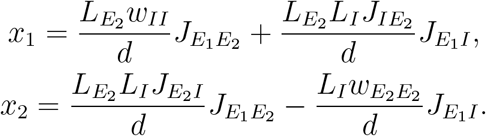

For sake of illustration assume that *d >* 0 (the same argument holds under reversal of signs). Then for 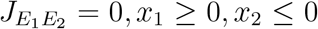 and 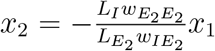, which is a lower bound for *x*_2_ in the region where *x*_1_ *>* 0. Notice with our assumption that *d* is positive, a similar argument letting 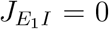 leads only to bounds on *x*_1_ ≤ 0, *x*_2_ ≥ 0 (since *w*_*II*_ ≤ 0). Thus we see how the *E*_2_ − *I* connectivity places additional constraints on the accessible values of *X* under modulation of connection strength (observe the white regions in Figure 6 for *x*_1_ ≥ 0, *x*_2_ ≤ 0 and *x*_1_ ≤ 0, *x*_2_ ≥ 0).

Different bounds will result from varying different parameters. For example, since the cellular gain, *L*_*α*_, is non-negative its minimum value is 0. Letting 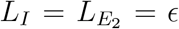 in equations 31 and 32 it can be shown that *x*_1_ *<* 0, *δ > x*_2_ ≥ 0 for some *δ* which depends on *ϵ* in the parameter regime used in Figure 5. In particular, the assumption of 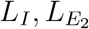 small leads to the terms in equations 31 and 32 with *w*’s dominating. These values would in turn cross the *J* -induced boundary, but do not violate our conclusions since *L*_*α*_ was assumed fixed in Figure 5.

**Figure S2:**
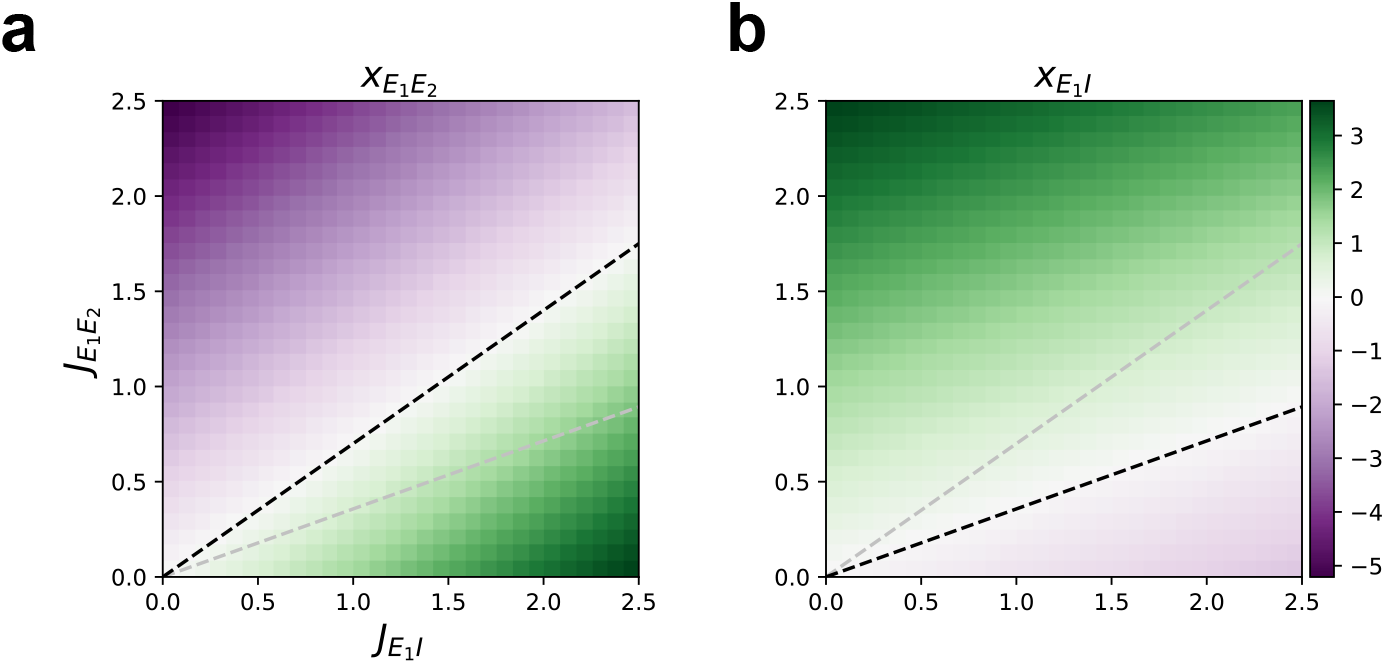
X components as a function of connectivity. **a**. 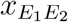 as a function of 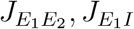. Black dashed line corresponds to 0 contour; gray dashed line is 0 contour of panel (b). **b**. Same as (a) for 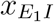; gray dashed line is now 0 contour of panel (a). Region bounded by dashed lines is where 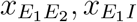 are jointly positive.

### *X* as paths through the network

As discussed above, in full generality 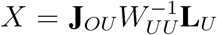 where *O* is the observed (readout) population, *U* the unobserved population. We recall that *W*_*UU*_ = −**1** + **L**_*U*_ **J**_*UU*_. Assuming boundedness of *W*_*UU*_ we have the expansion 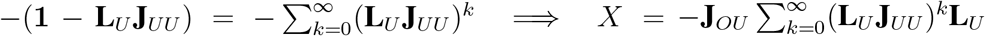. Since **L**_*U*_ **J**_*UU*_ are the effective connection strengths in the network, (**L**_*U*_ **J**_*UU*_)^*k*^ denotes the *k*^*th*^ step through the unobserved population [48], and consequently *X* is comprised of all paths through the unobserved population and their projections to the observed population (**J**_*OU*_). This expansion has been used before to decompose the structure of the response covariance in a neural network [56, 4].

The final panel in Figure 5 additionally derives from this expansion in the following way: consider the *ij* element of *X* where *i* is the index of a decoded unit and *j* is an undecoded unit. Then 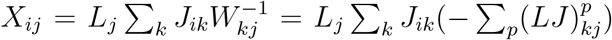. In the final sum the *kj*^*th*^ element is all the *p*-step ways to reach element *k* from element *j*, and in turn, the readout neuron *i*.

## References

[1] L. F. Abbott and P. Dayan. The effect of correlated variability on the accuracy of a population code. Neural computation, 11(1):91–101, 1999.

[2] B. B. Averbeck, P. E. Latham, and A. Pouget. Neural correlations, population coding and computation. Nature Reviews Neuroscience, 7(5):358–366, 2006.

[3] J. Beck, V. R. Bejjanki, and A. Pouget. Insights from a simple expression for linear fisher information in a recurrently connected population of spiking neurons. Neural computation, 23(6):1484–1502, 2011.

[4] H. Bos, A.-M. Oswald, and B. Doiron. Untangling stability and gain modulation in cortical circuits with multiple interneuron classes. bioRxiv, 2020.

[5] M. R. Cohen and J. H. Maunsell. Attention improves performance primarily by reducing interneuronal correlations. Nature neuroscience, 12(12):1594–1600, 2009.

[6] J. De La Rocha, B. Doiron, E. Shea-Brown, K. Josić, and A. Reyes. Correlation between neural spike trains increases with firing rate. Nature, 448(7155):802–806, 2007.

[7] B. Doiron, A. Litwin-Kumar, R. Rosenbaum, G. K. Ocker, and K. Josić. The mechanics of state-dependent neural correlations. Nature neuroscience, 19(3):383–393, 2016.

[8] A. S. Ecker, G. H. Denfield, M. Bethge, and A. S. Tolias. On the structure of neuronal population activity under fluctuations in attentional state. Journal of Neuroscience, 36(5):1775–1789, 2016.

[9] A. Fontanini and D. B. Katz. Behavioral states, network states, and sensory response variability. Journal of neurophysiology, 100(3):1160–1168, 2008.

[10] C. Gardiner. Stochastic methods, volume 4. springer Berlin, 2009.

[11] M. Getz, C. Huang, and B. Doiron. Understanding modulatory effects on cortical circuits through subpopulation coding. BMC Neuroscience, 2019. 28th Annual Computational Neuroscience Meeting: CNS*2019.

[12] M. Getz, C. Huang, and B. Doiron. Supopulation coding reveals a mechanism for improved information flow through cortical circuits. Cosyne Abstracts, Denver, CO, 2020.

[13] M. Getz, C. Huang, J. Dunworth, M. R. Cohen, and B. Doiron. Attentional modulation of neural covariability in a distributed circuit-based population model. Cosyne Abstracts, Denver, CO, 2018.

[14] L. M. Giocomo and M. E. Hasselmo. Neuromodulation by glutamate and acetylcholine can change circuit dynamics by regulating the relative influence of afferent input and excitatory feedback. Molecular neurobiology, 36(2):184–200, 2007.

[15] K. D. Harris and T. D. Mrsic-Flogel. Cortical connectivity and sensory coding. Nature, 503(7474):51–58, 2013.

[16] K. D. Harris and G. M. Shepherd. The neocortical circuit: themes and variations. Nature neuroscience, 18(2):170–181, 2015.

[17] K. D. Harris and A. Thiele. Cortical state and attention. Nature reviews neuroscience, 12(9):509–523, 2011.

[18] L. J. Herstel and C. J. Wierenga. Network control through coordinated inhibition. Current Opinion in Neurobiology, 67:34–41, 2021.

[19] C. Huang. Modulation of the dynamical state in cortical network models. Current opinion in neurobiology, 70:43–50, 2021.

[20] C. Huang, D. A. Ruff, R. Pyle, R. Rosenbaum, M. R. Cohen, and B. Doiron. Circuit models of low-dimensional shared variability in cortical networks. Neuron, 101(2):337–348, 2019.

[21] K. Josić, E. Shea-Brown, B. Doiron, and J. de la Rocha. Stimulus-dependent correlations and population codes. Neural computation, 21(10):2774–2804, 2009.

[22] M. Kafashan, A. Jaffe, S. N. Chettih, R. Nogueira, I. Arandia-Romero, C. D. Harvey R. Moreno-Bote, and J. Drugowitsch. Scaling of information in large neural populations reveals signatures of information-limiting correlations. bioRxiv, 2020.

[23] T. Kanashiro, G. K. Ocker, M. R. Cohen, and B. Doiron. Attentional modulation of neuronal variability in circuit models of cortex. Elife, 6:e23978, 2017.

[24] I. Kanitscheider, R. Coen-Cagli, and A. Pouget. Origin of information-limiting noise correlations. Proceedings of the National Academy of Sciences, 112(50):E6973–E6982, 2015.

[25] A. J. Keller, M. M. Roth, and M. Scanziani. Feedback generates a second receptive field in neurons of the visual cortex. Nature, pages 1–5, 2020.

[26] A. Kohn, R. Coen-Cagli, I. Kanitscheider, and A. Pouget. Correlations and neuronal population information. Annual review of neuroscience, 39:237–256, 2016.

[27] J. W. Krakauer, A. A. Ghazanfar, A. Gomez-Marin, M. A. MacIver, and D. Poeppel. Neuroscience needs behavior: correcting a reductionist bias. Neuron, 93(3):480–490, 2017.

[28] G. W. Lindsay, D. B. Rubin, and K. D. Miller. A simple circuit model of visual cortex explains neural and behavioral aspects of attention. bioRxiv, 2019.

[29] A. Litwin-Kumar and B. Doiron. Slow dynamics and high variability in balanced cortical networks with clustered connections. Nature neuroscience, 15(11):1498–1505, 2012.

[30] C. Ly, J. W. Middleton, and B. Doiron. Cellular and circuit mechanisms maintain low spike co-variability and enhance population coding in somatosensory cortex. Frontiers in Computational Neuroscience, 6:7, 2012.

[31] E. Marder. Neuromodulation of neuronal circuits: back to the future. Neuron, 76(1):1–11, 2012.

[32] E. Marder, T. O’Leary, and S. Shruti. Neuromodulation of circuits with variable parameters: single neurons and small circuits reveal principles of state-dependent and robust neuromodulation. Annual review of neuroscience, 37:329–346, 2014.

[33] J. C. Martinez-Trujillo and S. Treue. Feature-based attention increases the selectivity of population responses in primate visual cortex. Current biology, 14(9):744–751, 2004.

[34] C. J. McAdams and J. H. Maunsell. Effects of attention on orientation-tuning functions of single neurons in macaque cortical area v4. Journal of Neuroscience, 19(1):431–441, 1999.

[35] M. J. McGinley, M. Vinck, J. Reimer, R. Batista-Brito, E. Zagha, C. R. Cadwell, A. S. Tolias, J. A. Cardin, and D. A. McCormick. Waking state: rapid variations modulate neural and behavioral responses. Neuron, 87(6):1143–1161, 2015.

[36] J. S. Montijn, R. G. Liu, A. Aschner, A. Kohn, P. E. Latham, and A. Pouget. Strong information-limiting correlations in early visual areas. bioRxiv, page 842724, 2019.

[37] R. Moreno-Bote, J. Beck, I. Kanitscheider, X. Pitkow, P. Latham, and A. Pouget. Information-limiting correlations. Nature neuroscience, 17(10):1410–1417, 2014.

[38] F. Najafi, G. F. Elsayed, R. Cao, E. Pnevmatikakis, P. E. Latham, J. P. Cunningham, and A. K. Churchland. Excitatory and inhibitory subnetworks are equally selective during decisionmaking and emerge simultaneously during learning. Neuron, 105(1):165–179, 2020.

[39] D. H. Perkel and T. H. Bullock. Neural coding. Neurosciences Research Program Bulletin, 1968.

[40] M. I. Posner. Cognitive Neuroscience of Attention. Guilford Press, 2012.

[41] N. C. Rabinowitz, R. L. Goris, M. Cohen, and E. P. Simoncelli. Attention stabilizes the shared gain of v4 populations. Elife, 4:e08998, 2015.

[42] A. Renart and M. C. van Rossum. Transmission of population-coded information. Neural computation, 24(2):391–407, 2012.

[43] M. Rigotti, O. Barak, M. R. Warden, X.-J. Wang, N. D. Daw, E. K. Miller, and S. Fusi. The importance of mixed selectivity in complex cognitive tasks. Nature, 497(7451):585–590, 2013.

[44] D. B. Rubin, S. D. Van Hooser, and K. D. Miller. The stabilized supralinear network: A unifying circuit motif underlying multi-input integration in sensory cortex. Neuron, 85(2):402–417, 2015.

[45] D. A. Ruff and M. R. Cohen. Attention can either increase or decrease spike count correlations in visual cortex. Nature neuroscience, 17(11):1591–1597, 2014.

[46] P.-A. Salin and J. Bullier. Corticocortical connections in the visual system: structure and function. Physiological reviews, 75(1):107–154, 1995.

[47] S. Saxena and J. P. Cunningham. Towards the neural population doctrine. Current opinion in neurobiology, 55:103–111, 2019.

[48] A. C. Schwarze and M. A. Porter. Motifs for processes on networks. arXiv preprint 2007.07447, 2020.

[49] H. S. Seung and H. Sompolinsky. Simple models for reading neuronal population codes. Proceedings of the National Academy of Sciences, 90(22):10749–10753, 1993.

[50] G. M. Shepherd. Corticostriatal connectivity and its role in disease. Nature Reviews Neuroscience, 14(4):278–291, 2013.

[51] C. Stringer, M. Michaelos, D. Tsyboulski, S. E. Lindo, and M. Pachitariu. High-precision coding in visual cortex. Cell, 2021.

[52] C. Stringer, M. Pachitariu, N. Steinmetz, M. Carandini, and K. D. Harris. High-dimensional geometry of population responses in visual cortex. Nature, 571(7765):361–365, 2019.

[53] A. Thiele. Muscarinic signaling in the brain. Annual review of neuroscience, 36:271–294, 2013.

[54] A. Thiele and M. A. Bellgrove. Neuromodulation of attention. Neuron, 97(4):769–785, 2018.

[55] R. Tremblay, S. Lee, and B. Rudy. Gabaergic interneurons in the neocortex: from cellular properties to circuits. Neuron, 91(2):260–292, 2016.

[56] J. Trousdale, Y. Hu, E. Shea-Brown, and K. Josić. Impact of network structure and cellular response on spike time correlations. PLoS computational biology, 8(3):e1002408, 2012.

[57] R. S. Williamson and D. B. Polley. Parallel pathways for sound processing and functional connectivity among layer 5 and 6 auditory corticofugal neurons. Elife, 8:e42974, 2019.

